# Hypomineralized Enamel Alters Trigeminal Sensory Afferent Architecture, Transcriptome, and Dental Injury Responses

**DOI:** 10.64898/2026.09.10.750650

**Authors:** Hakan Karaaslan, Robert Bossong, Esther Wu, Lama Alabdulaaly, Min Son, Xuesong He, Isaac M. Chiu, Jennifer L. Gibbs, Felicitas B. Bidlack, Ozge Erdogan

**Affiliations:** Mineralized Tissue Biology and Bioengineering, ADA Forsyth Institute, Somerville, MA, United States; Department of Restorative Dentistry and Biomaterials Sciences, Harvard School of Dental Medicine, Boston, MA, United States; Maxillofacial Surgery and Diagnostic Sciences Department, College of Dentistry, King Saud bin Abdulaziz University for Health Sciences, Riyadh, Saudi Arabia; Department of Microbiology, ADA Forsyth Institute, Somerville, MA, United States; Department of Immunology, Blavatnik Institute, Harvard School of Medicine, Boston, MA, United States; Department of Endodontics, Department of Translational Dental Medicine, Boston University Henry M. Goldman School of Dental Medicine, Boston, MA, United States

## Abstract

At barrier tissues such as the skin and gut, sensory neurons interface with environmental stimuli and coordinate with neighboring epithelial and immune cells to detect tissue perturbations. In teeth, however, the sensory dentin-pulp complex is insulated from the oral environment by highly mineralized enamel. Although dentin-pulp responses to severe injury have been studied in models with direct pulp exposure, it remains unclear whether enamel barrier dysfunction alone alters pulpal and neuronal homeostasis. Using a kallikrein-related peptidase 4 knockout (KLK4 KO) mouse model of enamel hypomineralization, we demonstrated that defective enamel induced structural, molecular, and transcriptional responses in the dental pulp and in the trigeminal system innervating teeth, despite the absence of direct pulp exposure to the oral cavity. Hypomineralized molars exhibited increased reactionary dentin formation accompanied by retraction of sensory afferents from the dentin-pulp junction. Consistent with these structural findings, trigeminal ganglia of KLK4 KO mice displayed upregulation of genes associated with cytoskeletal remodeling and stimulus response pathways. Despite increased bacterial burden and biofilm accumulation on the enamel surface, enamel hypomineralization did not induce substantial innate immune responses in the pulp. In addition, following severe dental pulp injury, teeth with hypomineralized enamel exhibited reduced sensory afferent loss and tissue damage. Together, these findings reveal that enamel integrity functions as a critical regulator of dentin-pulp homeostasis and that barrier dysfunction alone can precondition tissue responses to subsequent injury.

## INTRODUCTION

Enamel is a highly specialized mineralized barrier that forms the outermost layer of the tooth crown. Enamel-producing cells, ameloblasts, die as the tooth erupts, leaving an acellular mineralized matrix that covers and protects the dental pulp and dentin (1). Dental pulp is the sensory organ of the tooth and is densely innervated by sensory neurons of the trigeminal ganglion (2). Dentin lies between the enamel and the pulp and acts as an additional mineralized barrier. Unlike most barrier tissues, where sensory afferents lie close to epithelial surfaces and readily encounter environmental stimuli and microbes (3, 4), sensory afferents within the dental pulp are spatially segregated from the external environment by these highly specialized mineralized barriers.

Dentin, produced by odontoblasts, is a cellular, tubular mineralized tissue that can respond to external and internal stimuli throughout an individual’s lifespan (5). The dental pulp and sensory afferents are in direct contact with dentin and odontoblasts, and together they form a sensory protective interface called the dentin-pulp complex (6, 7). Any insult that reaches dentin is considered to expose the dental pulp and its innervation, and this complex contributes to physiological functions such as sensing the external environment, defense, and repair (8, 9). Similarly, any defect that compromises the barrier function of enamel can potentially greatly influence how dentin and dental pulp perceive and respond to physical, chemical, and microbial stimuli. However, the possibility that defects in the enamel barrier may also influence the dental pulp sensory organ has received little attention based on the anatomical distance between enamel and pulp.

Molar hypomineralization (MH) is a common developmental defect of enamel with sporadic, localized hypomineralization, presenting as demarcated opacities. These opaque patches of defective enamel have lower mineral density, higher protein content, and increased porosity (10–12). Clinical studies suggest that structural changes in enamel, as seen in MH, can increase dental hypersensitivity and can even lead to failure of local anesthesia during dental procedures (13–16). Clinical symptoms suggest sensory alterations in the dental pulp. Studies of extracted teeth from patients with molar hypomineralization further demonstrate altered expression of sensory neuronal markers, including TRPV1, in the dental pulp (17, 18). However, the nature and extent of these alterations, and how impairment of the enamel barrier affects dental pulp sensory afferents and other dentin and pulp cells, are not known.

To investigate the primary impact of hypomineralized enamel on pulpal sensory dynamics, we utilized a kallikrein-related peptidase 4 knockout (KLK4 KO) mouse model exhibiting a hypomineralized enamel phenotype (19). KLK4 is a key protease responsible for removing enamel matrix proteins during enamel maturation; loss of this enzyme results in a hypomineralized enamel resembling the demarcated opacities observed in MH (20). Second, this model also allowed us to recapitulate a commonly observed disease progression, in which structurally compromised enamel lesions can easily progress to deep caries cavities that expose the dental pulp. Combining the KLK4 KO model and a previously established model of pulp exposure, we studied how the dental pulp responds to more severe injuries in the presence of an existing enamel barrier defect.

We hypothesize that, even in the absence of active caries or direct bacterial invasion of the dentin, measurable sensory and immune alterations within the dental pulp and the trigeminal system would result from increased exposure to both bacterial and non-bacterial stimuli. These alterations can then potentially prime the dental pulp and change its response to subsequent insults. We investigated the structural, molecular, and transcriptional responses to hypomineralized enamel in the dental pulp and trigeminal system. Understanding how enamel functions as a barrier to protect the sensory organ of the tooth from these stimuli will ultimately lead to improved treatment strategies for patients affected by non-carious lesions and developmental defects of the tooth, such as MH.

## RESULTS

To test our hypothesis, we compared KLK4 KO animals with hypomineralized enamel to WT animals at baseline, without any treatment. Compared with the hard and shiny enamel of WT animals (Figure 1B), KLK4 KO animals have soft and opaque enamel (Figure 1C) resembling human hypomineralized enamel. Hypomineralized enamel has lower mineral density (Figure 1B, represented with a darker shade on microCT images) when compared to the healthy enamel of WT animals (Figure 1D), yet it still covers most of the dentin.

**Figure 1.**
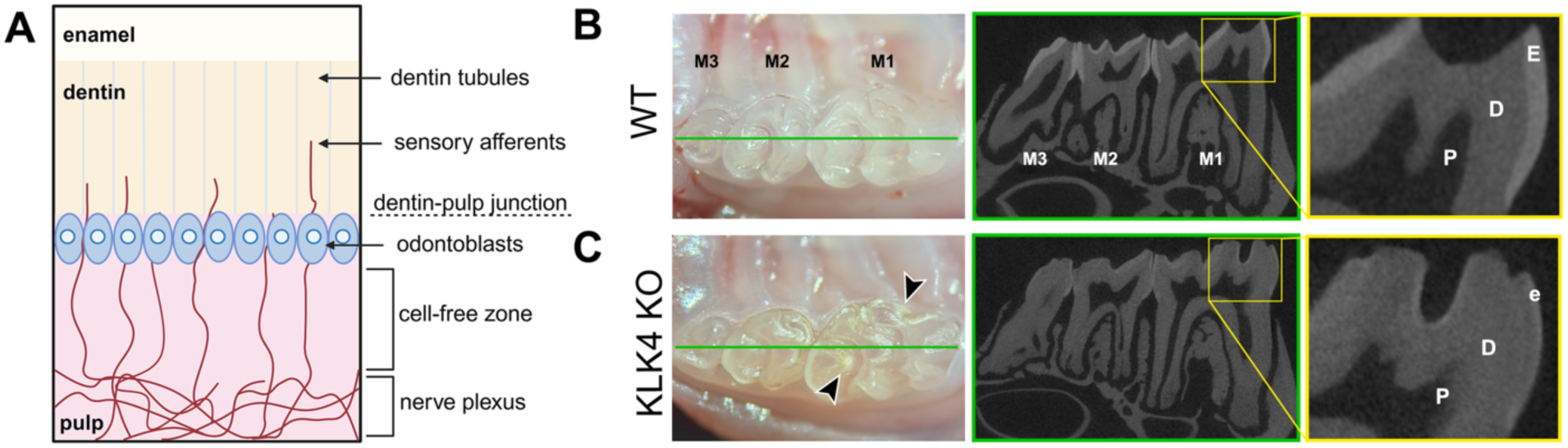
Spatial description of enamel, dentin, and the pulp, and description of the KLK4 KO hypomineralized enamel model. **(A)** Schematic showing the spatial relation of enamel, dentin, and pulp, as well as the odontoblast and sensory afferents. Intraoral photographs and microCT images of **(B)** WT and **(C)** KLK4 KO molar teeth. While still covering the underlying dentin and pulp, KLK4 KO enamel has lower mineral density (less white on microCT images), similar to hypomineralized human enamel. Similar to the demarcated opacities seen in patients with Molar Hypomineralization, the hypomineralized enamel of KLK4 KO animals looks opaque and chalky (black arrow heads), compared to the shiny surface of the healthy enamel of WT animals. E: Enamel, e: hypomineralized enamel, D: Dentin, P: Pulp, M1: Maxillary 1^st^ molar, M2: 2^nd^ molar, M3: 3^rd^ molar tooth.

### Hypomineralized enamel leads to increased reactionary dentin formation and retraction of sensory afferents from dentin

Odontoblasts are located at the dentin-pulp junction (Figure 1A), and they produce reactionary dentin when they receive insult or injury. To assess odontoblast response to hypomineralized enamel, we compared reactionary dentin formation in the molar teeth of WT and KLK4 KO animals, using the irregular tubule structure on the DAPI-labeled phase contrast images. KLK4 KO molar teeth with hypomineralized enamel showed a significant increase in reactionary dentin formation when compared to WT teeth (Fig. 2A). Quantification of the reactionary dentin area (normalized to the total pulp area) revealed a significant elevation in reactionary dentin formation across all cusps when analyzed combined (p<0.05) (Fig. 2B). When analyzed for individual cusps, the only significant difference was observed at the middle cusp. While the mesial and distal cusps still showed similar trends, they were not statistically significant (p>0.05).

**Figure 2.**
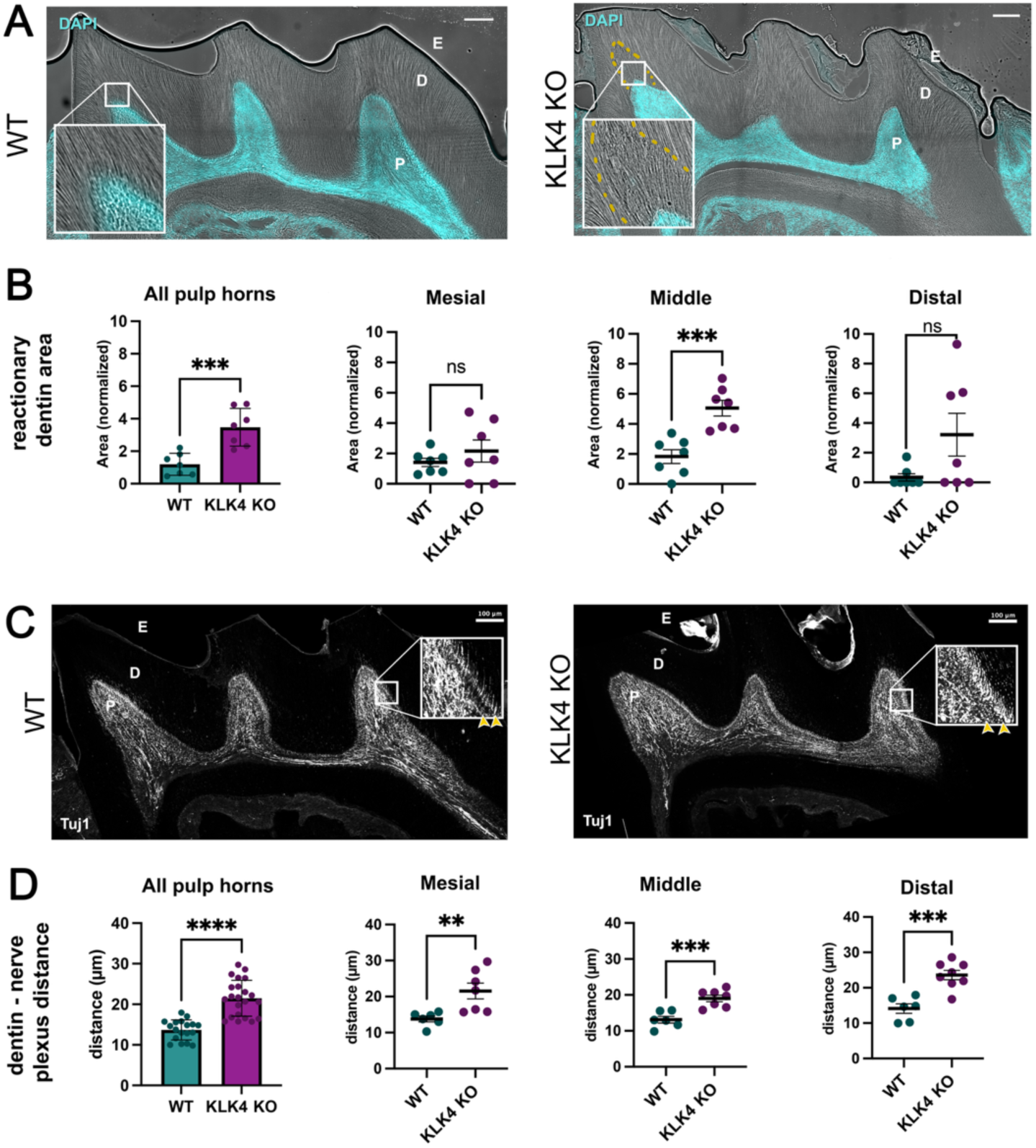
Hypomineralized enamel leads to increased reactionary dentin formation and sensory afferent retraction from the dentin-pulp junction. **(A)** Representative phase contrast images showing reactionary dentin formation in WT and KLK4 KO first molars. The yellow dashed line delineates the reactionary dentin with irregular tubule formation in the KLK4 KO molar, and cyan (DAPI) shows all cells in the dental pulp. **(B)** The reactionary dentin area is significantly larger in the KLK4 KO molars, when compared to WT across all pulp horns. When each pulp horn is evaluated separately between WT and KLK4 KO, the difference is only significant in the middle pulp horn. **(C)** Representative Tuj1-stained immunofluorescence images of the first molars of WT and KLK4 KO animals, showing the sensory afferents in the dental pulp. Sensory afferent plexus (Raschkow’s Plexus) is withdrawn away from the dentin underneath hypomineralized enamel. Representative reference points for measuring the distance between dentin and the sensory plexus are indicated with yellow arrows. **(D)** The distance between dentin and the sensory afferent plexus is significantly increased in the KLK4 KO molars, when compared to WTs across all pulp horns. Comparison of all pulp horns separately also indicates a significant increase in this distance between WT and KLK4 KO. E: Enamel, D: Dentin, P: Pulp, Cyan: DAPI. Scale bars: 100 μm. Mean ± SEM; Two-way ANOVA with Tukey’s or Sidak’s multiple comparisons test; **p < .01, ***p < .001, ****p < .0001, ns = not significant.

To assess the structural organization of sensory afferents in teeth with hypomineralized enamel, we performed βIII-tubulin (Tuj1) immunostaining to label nerve fibers in molar tooth sections from WT and KLK KO mice. Compared to healthy molar teeth in WT animals, molar teeth of KLK4 KO animals with hypomineralized enamel displayed a significant increase in the distance between the nerve plexus formed by Tuj1⁺ nerve terminals and the dentin-pulp junction (Fig. 2C), indicating a retraction of sensory afferents from the dentin-pulp junction (Fig. 2C-D). This retraction was observed consistently across all three cusps and was statistically significant when analyzed individually and combined (Fig. 2D). These findings suggest that hypomineralized can lead to remodeling of the sensory architecture and increased reactionary dentinogenesis.

### Sensory neuron activation and sensory neuron stress marker expression are increased in trigeminal nuclei and trigeminal ganglia, respectively, due to hypomineralized enamel

To determine whether hypomineralized enamel leads to an alteration in sensory neuron activation and whether it affects sensory neurons in the dental pulp, we assessed the expression of the immediate early gene, c-Fos, in the trigeminal nuclei, which is a marker for neuronal activity, and ATF-3, a transcription factor in the trigeminal ganglia, which is a marker of neuronal injury. First, we quantified and compared c-Fos expression in the trigeminal nuclei. The percentage area of c-Fos-positive cells in the trigeminal nuclei of KLK4 KO mice was significantly higher when compared with WT controls (Fig. 3B-C; p < 0.05), indicating heightened neuronal activation. We next examined the trigeminal ganglia for evidence of neuronal injury. We found that ATF3 expression was significantly higher in the KLK4 KO trigeminal ganglia than in WT (Fig. 3D-E; p < 0.05). Together, these results suggest that hypomineralized enamel leads to enhanced neuronal activation and increased peripheral injury responses in the trigeminal system, indicating an overall sensitized state in animals with hypomineralized enamel.

**Figure 3.**
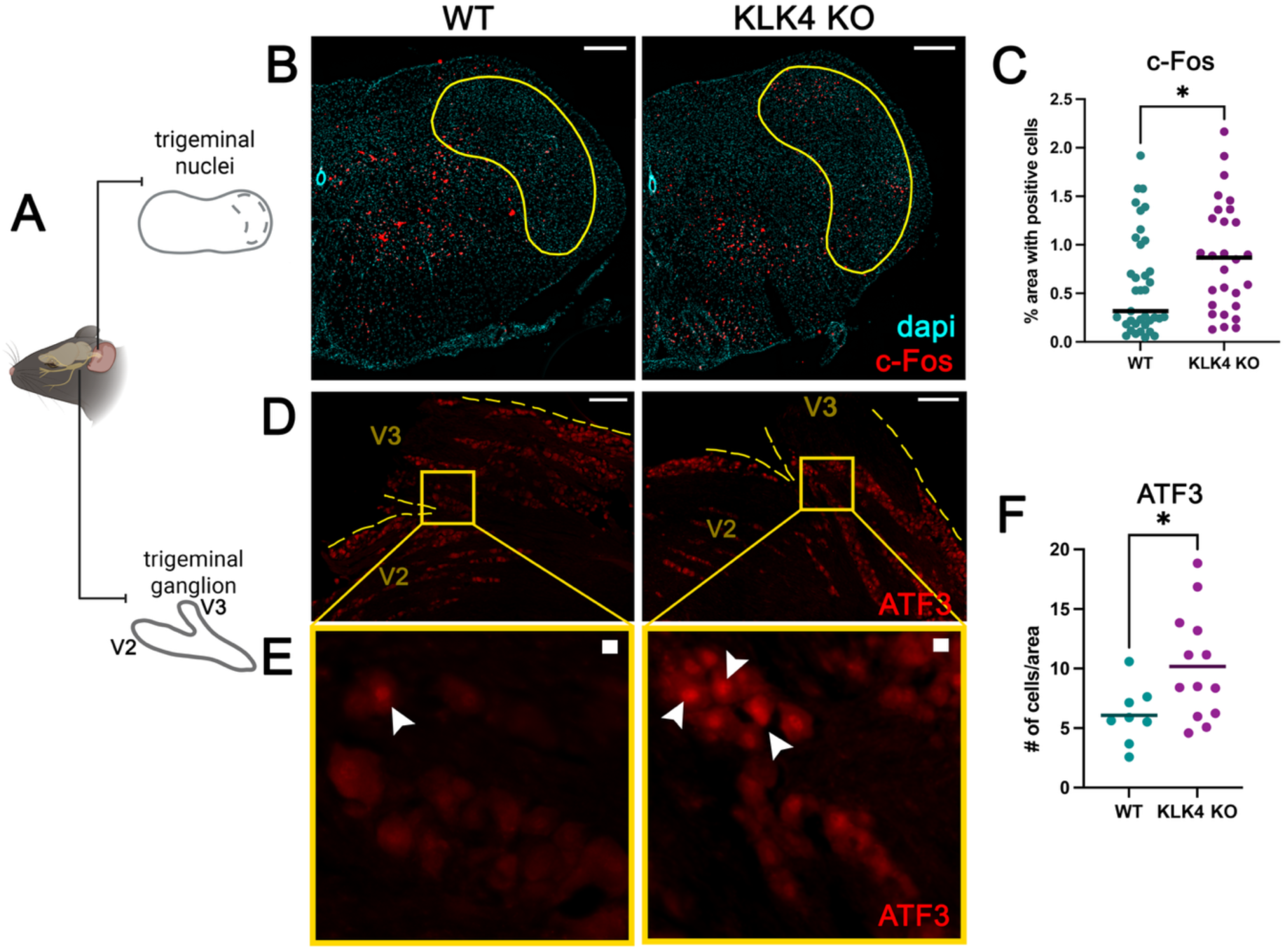
Sensory neurons are activated and exhibit increased expression of neuronal stress markers beneath hypomineralized enamel. **(A)** Schematic representation of the trigeminal ganglion and brainstem containing the trigeminal nuclei. **(B)** Representative images of coronal sections from the trigeminal nucleus caudalis and interpolaris with c-FOS (red) and DAPI (cyan) of KLK4 KO and WT animals. Scale bar = 200 μm. **(C)** The area with c-Fos-positive cells is significantly increased in the KLK4 KO animals when compared to WT. **(D)** Representative low-magnification images of the trigeminal ganglion of WT and KLK4 KO animals with ATF3 (red). The intersection area between V3 and V2 divisions can be seen (scale: 200 μm). **(E)** Arrowheads denote ATF3-positive neurons in the high-magnification images (scale = 10 μm). **(F)** The number of ATF3-positive cells per unit area was significantly higher in the KLK4 KO animals, when compared to WTs. Quantification of ATF3 was performed only in the region containing neurons innervating the teeth. Mean ± SEM; each dot represents an individual animal. Unpaired two-tailed t-tests; *p < .05.

### Trigeminal ganglia exhibit distinct transcriptional signatures in mice with hypomineralized enamel

To investigate the molecular consequences of hypomineralized enamel on sensory neurons in the teeth, we performed differential gene expression analysis using bulk RNA sequencing of trigeminal ganglia from WT and KLK4-KO mice. Among the most upregulated genes in KLK4 KO animals were *Stmn1*, a microtubule-destabilizing protein involved in axonal growth and regeneration, *Snhg3*, a long noncoding RNA implicated in transcriptional regulation and cellular stress responses, and *Nudc,* a protein involved in cell division and neuronal migration. Conversely, several genes were markedly downregulated in KLK4 KO mice, including *Abhd1*, an enzyme linked to lipid metabolism and mitochondrial function, *Cyp2j13*, a member of the CyP 450 family involved in lipid mediator synthesis, and *Pomc*, a neuropeptide precursor regulating stress responses, pain modulation, and neuroendocrine signaling (Fig. 4B).

**Figure 4.**
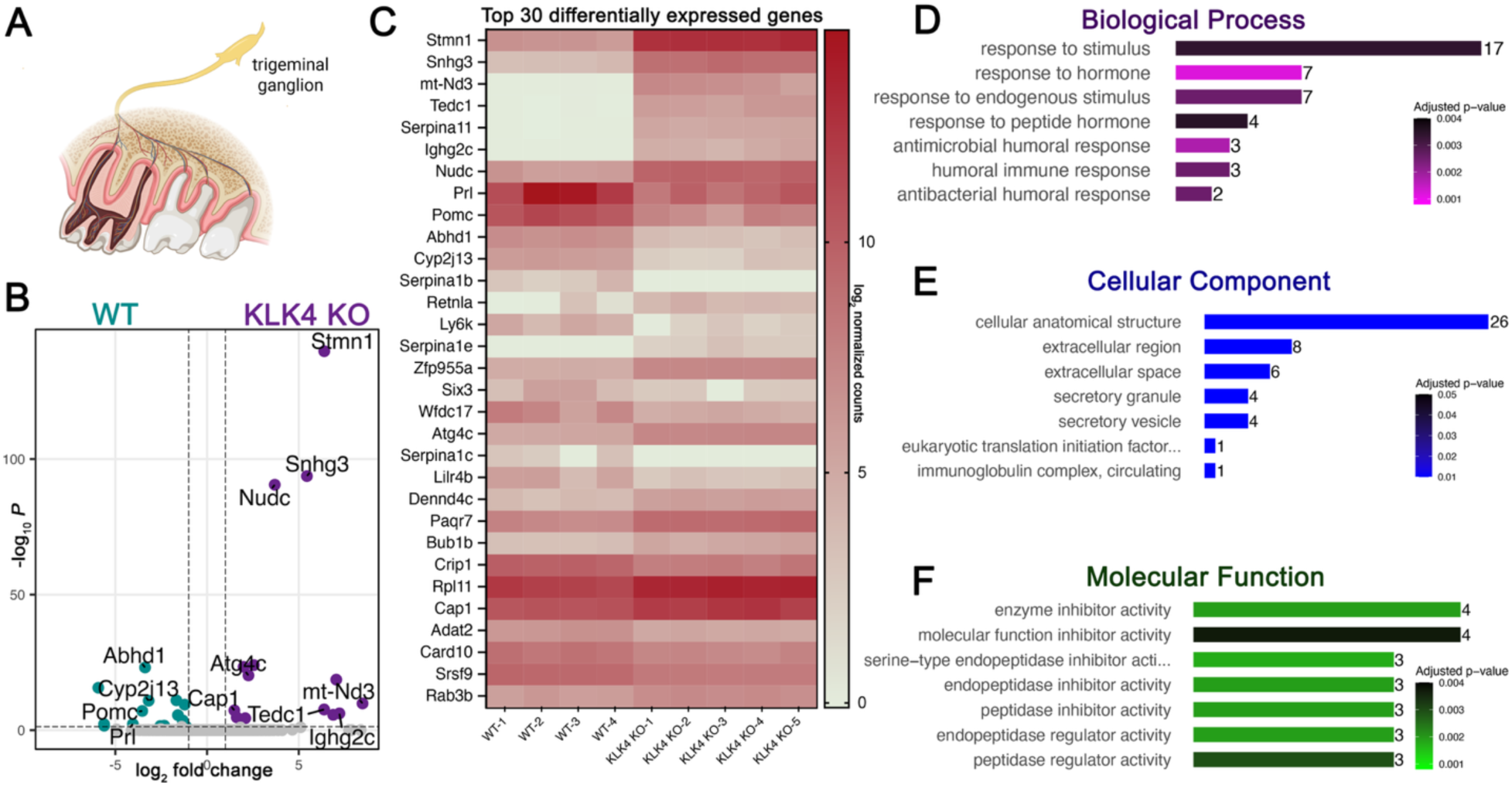
Transcriptomic analysis of the trigeminal ganglion of WT and KLK4 KO animals. **(A)** A schematic of the trigeminal ganglion, where the cell bodies of sensory neurons innervating the dental pulp reside, was collected and analyzed using bulk RNA sequencing. **(B)** Differentially expressed genes in KLK4 KO (purple) and WT (green) animals are shown on the volcano plot. Key genes are labeled. **(C)** The heatmap shows the top 30 differentially expressed genes in WT and KLK4 KOs, ranked by adjusted *p*-value; values are plotted on a log_2_ scale after normalization. **(D-F)** GO Enrichment Analysis shows the top seven differentially expressed groups based on the number of genes in KLK4 KO animals that are significant in **(D)** biological processes **(E)** cellular components, and **(F)** molecular function categories. The top groups in each category with the most genes are: response to stimulus (GO:0050896), cellular anatomical structure (GO:0110165), and enzyme inhibitor activity (GO:0004857), respectively. Bars indicate the number of genes in each category, and color shades indicate adjusted *p-*values.

A heatmap of the top 30 differentially expressed genes revealed clear clustering of WT and KLK4 KO samples, with distinct gene expression patterns in the two groups (Fig. 4C). Gene Ontology enrichment analysis showed that altered genes were involved in *biological processes* such as *“response to stimulus,” “response to hormone,”* and *“antimicrobial humoral response”* (Fig. 4D). In the *cellular component* category, enriched terms included *“cellular anatomical structure,” “extracellular region,”* and *“secretory vesicle”* (Fig. 4E). *Molecular function* enrichment highlighted *“enzyme inhibitor activity,” “serine-type endopeptidase inhibitor activity,”* and related regulatory roles (Fig. 4F). Together, these results indicate that molar hypomineralization can drive transcriptional reprogramming of the sensory neurons of the teeth which can be detected in the trigeminal ganglion, characterized by changes in genes regulating neuronal structural dynamics, stress signaling, and potential lipid metabolism, and immune response pathways.

### Myeloid cell populations in the dental pulp are comparable in WT and KLK4 KO animals

Resident macrophages constitute the major resident immune cell population of the dental pulp. To determine whether hypomineralized enamel alters the baseline immune composition of the dental pulp, we compared myeloid immune cell populations of the dental pulp between KLK4 KO and wild-type mice using flow cytometry. The proportion of CD45⁺ immune cells among all live pulp cells was similar between the two groups (p > .05), accounting for approximately 5-7% of total live cells. Within the CD45⁺ compartment, the majority of the immune cells were CD11b+ myeloid cells, and the frequency of CD11b⁺ myeloid cells did not differ significantly between WT and KLK4 KO mice. Among CD11b⁺ myeloid cells, the majority were F4/80⁺ macrophages, and their proportions remained comparable between WT and KLK4 KO animals. Further subclassification of macrophages by CX3CR1 expression revealed no significant differences in the proportions of CX3CR1⁺ and CX3CR1⁻ subsets. A small population of F4/80⁻ myeloid cells was also present at similar percentages in both genotypes. Additionally, the percentages of Ly6G⁺ neutrophils within the CD11b⁺F4/80⁻ population were equivalent between groups, suggesting that neutrophil representation in the healthy pulp is unaffected by the enamel hypomineralizations (Fig. 5A-C). Together, these data indicate that hypomineralized enamel does not significantly impact the baseline composition of myeloid immune populations, including macrophages and neutrophils, in the dental pulp.

**Figure 5.**
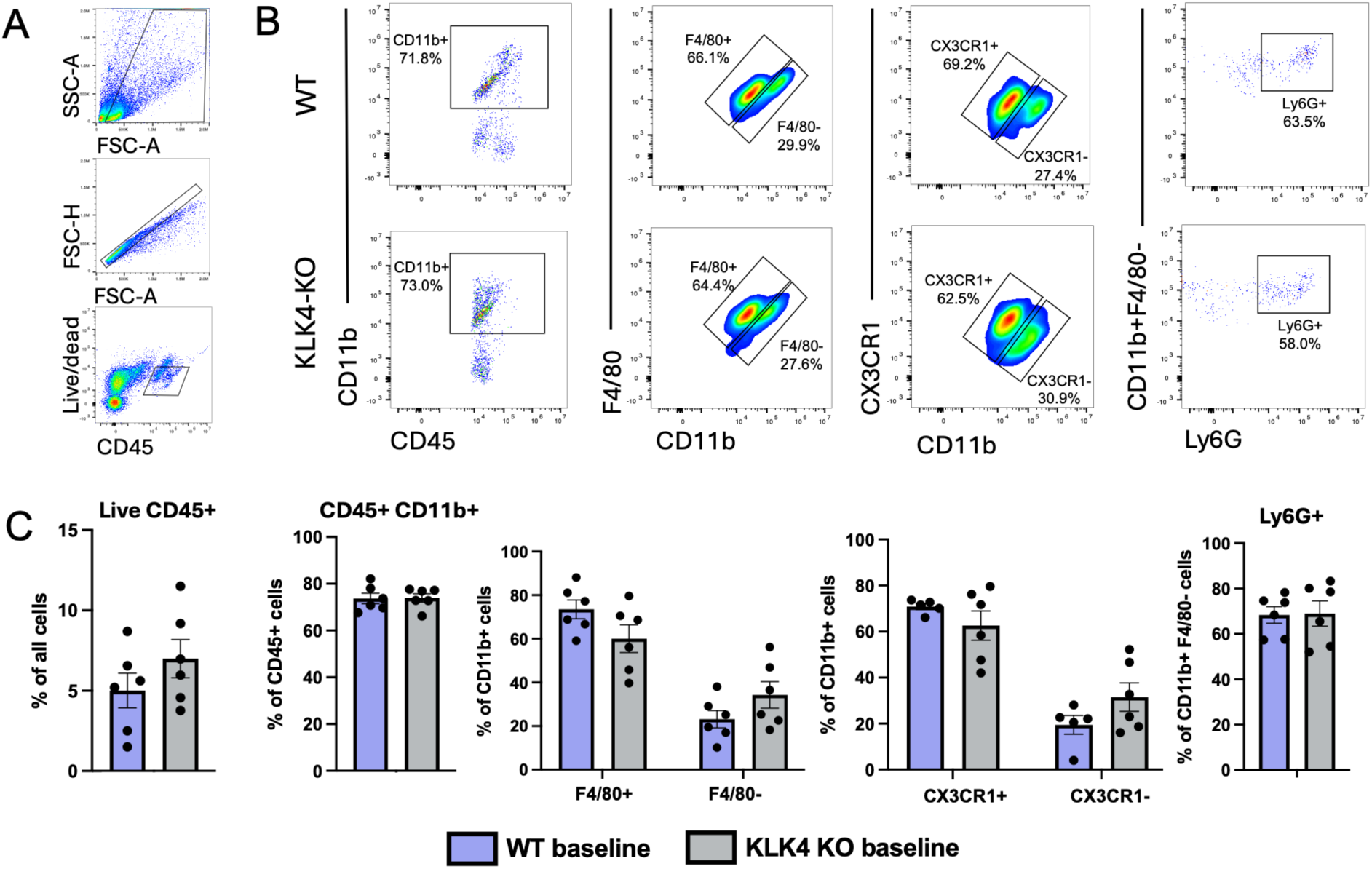
Myeloid cell populations are comparable in WT and KLK4 KO dental pulp under homeostatic conditions. **(A)** Representative flow cytometry (FC) plots showing the initial gating strategy for live, single, CD45⁺ immune cells. **(B)** Representative FC plots from WT and KLK4 KO dental pulp showing: CD45⁺CD11b⁺ myeloid cells, F4/80⁺ macrophages among CD11b⁺ cells, CX3CR1⁺ macrophages among CD11b⁺ cells, and Ly6G⁺ neutrophils among CD11b⁺F4/80⁻ cells. **(C)** Quantification of immune cell subsets showing similar proportions of CD45⁺ immune cells, CD11b⁺ myeloid cells, macrophages (F4/80⁺) and F4/80^-^ myeloid cell subset, CX3CR1⁺ macrophages and CX3CR1⁻ myeloid cell subset, and CD11b⁺F4/80⁻ Ly6G⁺ neutrophils between WT and KLK4 KO mice. Data are shown as mean ± SEM, each dot representing an individual biological replicate. Unpaired t-test or two-way ANOVA with Sidak’s or Tukey’s multiple comparisons tests were performed.

### Hypomineralized enamel is associated with a marked increase in oral bacterial biomass, while community composition is preserved

16S rRNA sequencing revealed that the overall taxonomic composition of the oral microbiome was similar between WT and KLK4 KO mice. At both the genus and species levels, relative abundance profiles were broadly conserved, with no major shifts in dominant taxa (Fig. 6A-B). Alpha diversity analysis showed comparable species richness between groups, with only a modest trend toward reduced richness in samples from KLK4 KO animals (Fig. 6C). Beta diversity analysis likewise demonstrated overlapping community clustering, indicating minimal compositional divergence between genotypes (Fig. 6D)

**Figure 6.**
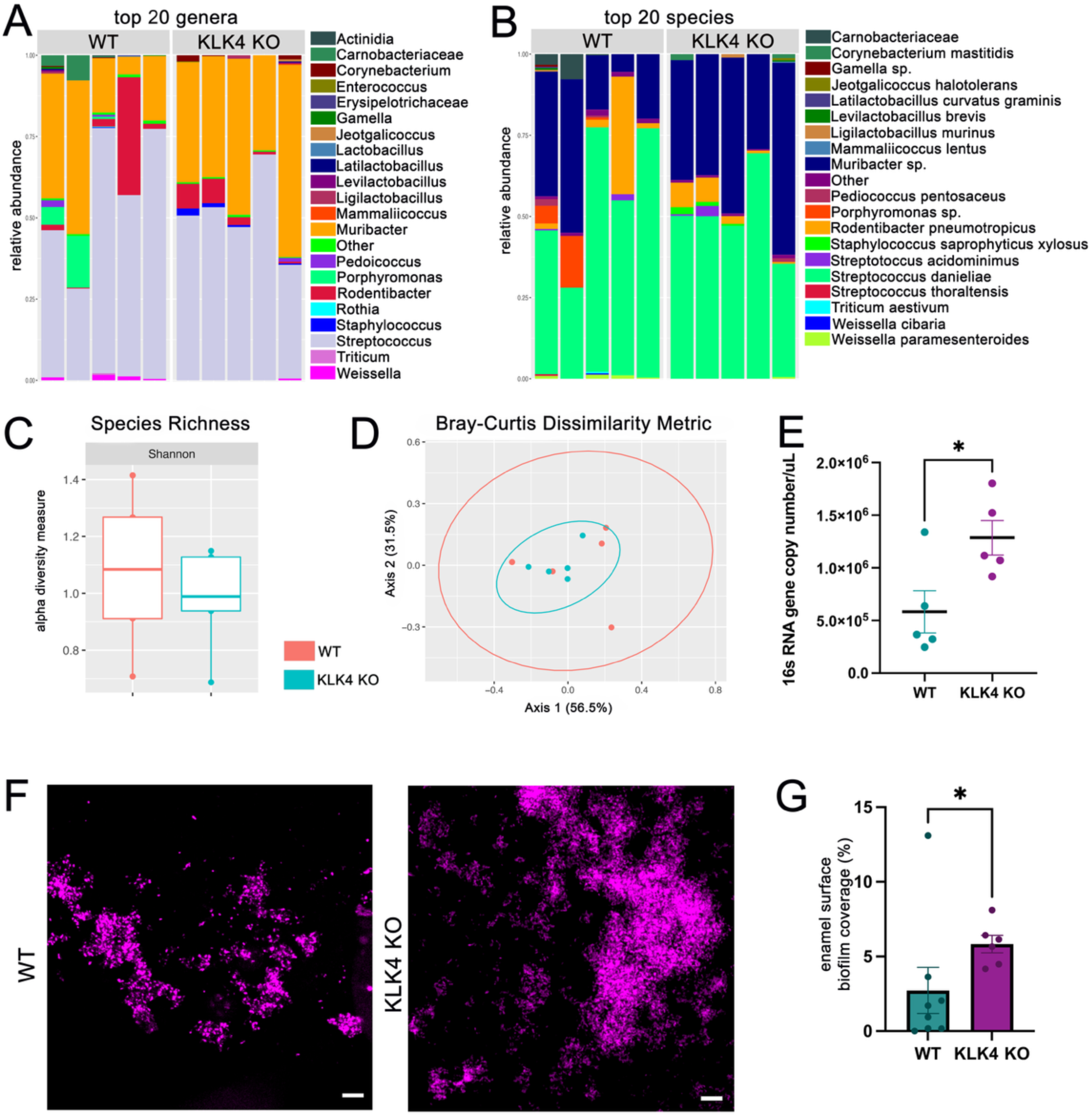
While oral microbiome composition is not altered, oral bacterial load is significantly higher in animals with hypomineralized enamel. **(A)** Relative abundance of the top 20 bacterial genera in the oral cavity of WT and KLK4 KO animals. **(B)** Relative abundance of the top 20 bacterial species in the oral cavity of WT and KLK4 KO animals. **(C)** Alpha diversity within groups is represented by Shannon alpha diversity values, and the **(D)** Bray-Curtis dissimilarity metric is used to assess beta diversity between the oral microbiomes of WT and KLK4 KO animals and was plotted by the NMDS. **(E)** The total oral bacterial load measured by 16s RNA gene copy number/μL was significantly higher in the KLK4 KO animals compared to the WT animals. **(F)** Representative images of in vitro grown 2-day Streptococcus mutans biofilm on extracted WT and KLK4 KO incisors. **(G)** Total biofilm coverage on the enamel surface was significantly higher in the KLK4 KO animals with hypomineralized enamel, when compared to WTs. Unpaired two-tailed t-tests; *P < .05. Scale bars: = 10 μm.

Despite the similarity in community structure, KLK4 KO mice exhibited significantly higher bacterial burden, as indicated by increased 16S rRNA gene copy numbers (Fig. 6E; p < .05). Consistent with this result, confocal imaging using FISH labelled *ex vivo* biofilm on enamel surface revealed greater surface coverage by *Streptococcus mutans* in KLK4 KO animals compared with WTs (Fig. 6F-G; p < .05). These findings suggest that while hypomineralized enamel does not substantially alter the overall composition of the oral microbiome, it promotes greater microbial accumulation and biofilm formation on enamel surfaces.

### Pulp tissue damage is reduced in the *primed* dental pulp beneath hypomineralized enamel after pulp injury

Based on our findings on the molecular and structural responses of the dental pulp to hypomineralized enamel, we asked whether the dental pulp beneath the hypomineralized enamel is *primed* for altered responses to other insults. To test this, we created dental pulp injuries in molar teeth in KLK4 KO and WT animals (experimental timeline is shown in Fig. 7I). At day three following the pulp injury, molar teeth from KLK4 KO animals showed significantly less sensory afferent loss when compared with WT animals (Fig. 7A, B; p < .05). Moreover, CD31 staining for endothelial cells showed a significant increase in vascular length per pulp area in the dental pulp from KLK4 KO animals compared to WT animals (Fig. 7C-D; p < .01).

**Figure 7.**
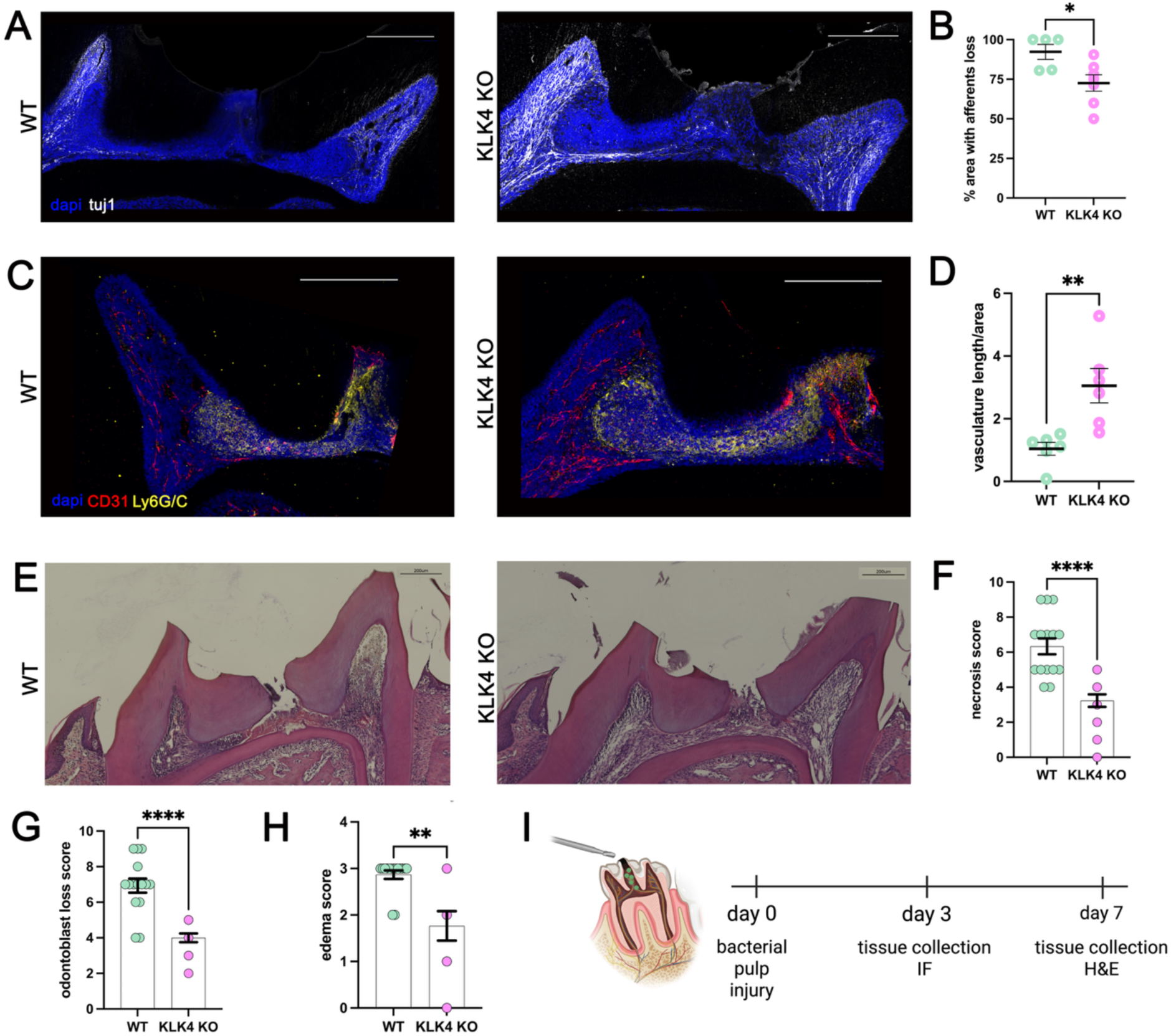
Pulp tissue damage is less in the *primed* dental pulp beneath hypomineralized enamel after pulp injury. **(A)** Representative Tuj1-stained immunofluorescence (IF) images of WT and KLK4 KO molars 3 days after pulp injury, showing sensory afferent loss inside the dental pulp. **(B)** When compared to WT, the area of the afferent loss was significantly less in the KLK4 KO. **(C)** Representative images of CD31 (endothelial cells) and Ly6G/C (neutrophils/monocytes) in mesial and mid pulp horns of molar teeth 3 days after pulp injury in WT and KLK4 KO animals. **(D)** The vasculature of the dental pulp was significantly denser after injury in the KLK4 KO when compared to WTs. **(E)** Representative images of hematoxylin and eosin-stained sections, 7 days after pulpal injury. When compared to WTs, KLK4 KO molars showed significantly less **(F)** pulp necrosis, **(G)** odontoblast loss, and **(H)** edema, **(I)** Timeline of the experimental procedures. Scale bars: 200 μm. Unpaired two-tailed t-tests; *P < .05, **P < .01, ****P < .0001. blue: DAPI.

By day seven, there was less tissue necrosis in the dental pulp in KLK4 KO animals compared to WTs (Fig. 7E), exhibiting significantly lower necrosis scores (Fig. 7F; p < .0001), reduced odontoblast loss (Fig. 7G; p < .0001), and decreased edema (Fig. 7H; p < .01) in the dental pulp of KLK4 KO animals compared with WTs. These findings suggest that dental pulp beneath hypomineralized enamel is in a primed state that protects it from greater injury.

## DISCUSSION

Enamel is a cell-free extracellular mineralized matrix and, in contrast to dentin and bone, is of epithelial origin (30, 31). Consequently, unlike other epithelial barriers, enamel lacks the capacity for adaptive cellular responses to insults (32). When the enamel barrier is compromised, cells in the dentin-pulp complex that are separated from enamel by the dentin layer respond. Even when the primary defect is strictly confined to enamel, the biological response is mediated the odontoblasts, sensory afferents and other dental pulp cells that reside in the underlying dentin-pulp complex (33). Since the dentin-pulp complex cannot repair enamel directly; instead, it responds by attempting to avoid or mitigate the impact of the stimulus through biological adaptation, such as the formation of reactionary dentin (34).

Odontoblasts, positioned at the dentin-pulp junction, form the first cellular barrier responding to any stimuli from the oral cavity (35). In response to mild or chronic stimuli such as early stage caries, and dentin defects, surviving odontoblasts increase secretory activity and deposit reactionary dentin (36–38). In line with these findings, molars with hypomineralized enamel in our model exhibited increased reactionary dentin formation, indicating increased odontoblast activity due to receiving augmented stimuli from the oral cavity. Prior studies have largely attributed this response to bacterial products or biomolecules released during dentin demineralization (39–42). In contrast, our findings suggest that even in the absence of caries, enamel defects can initiate odontoblast responses, although the precise molecular triggers remain unclear.

Sensory afferents are also located at the dentin pulp-junction, where they extend into the dentinal tubules, entangled with the odontoblast processes (43–45). We observed that in the molar teeth of KLK4 KO animals with hypomineralized enamel, the plexus formed by sensory afferents was retracted away from the dentin-pulp junction. This pattern resembles responses to chronic dentin injuries prepared experimentally to induce pulpal inflammation (46). In line with these structural findings, our trigeminal transcriptomic data showed increased expression of genes related to cytoskeletal remodeling and enrichment for pathways such as response to stimulus, cellular modeling, and antimicrobial activity. We also observed increased markers of neuronal activation (c-Fos) and injury (ATF3) in the trigeminal system of KLK4 KO animals. These responses are consistent with prior findings are similar to previous studies that showed increased c-Fos expression in the trigeminal nuclei when teeth were exposed to noxious cold, as well as bacterial stimuli (47), and increased ATF3 expression when the dental pulp was injured (48, 49). All these findings suggest that hypomineralized enamel, even when macroscopically intact and located distant, is sufficient to induce sustained sensory responses in the dentin-pulp complex at the molecular, cellular, and structural levels.

We hypothesize that the altered physicochemical properties of hypomineralized enamel, specifically increased porosity, greater surface roughness, and reduced mineral content, (10, 50–52) modify the transmission of environmental and bacterial stimuli to the dentin-pulp complex. Healthy enamel, even though it is ninety-five percent mineral by weight, is semi-permeable (53). Hypomineralized enamel, which has greater water content due to increased porosity, can therefore alter the sensory environment of odontoblasts, sensory afferents, and other pulp cells via altered fluid dynamics through the dentinal tubules (52, 54, 55). These longstanding changes in sensory input may contribute to the odontoblast and sensory neuronal alterations observed in our model, in addition to mechanical stress due to softer enamel. Additionally, the enhanced biofilm accumulation observed on hypomineralized enamel surfaces, even in the absence of compositional shifts in the microbiome, may increase exposure to microbial metabolites, eliciting a response in the dentin-pulp complex (56). Further studies are needed to identify the causal drivers of the odontoblast and sensory neuron responses due to enamel defects.

Despite the increased bacterial burden, we did not observe significant immune cell recruitment in the dental pulp of KLK4 KO animals. There were no differences in myeloid cell populations, specifically in F4/80⁺macrophage and Ly6G⁺neutrophil subsets, between WT and KLK4-KO animals. This contrasts with classical bacterial pulpitis models, where significant immune cell recruitment, especially neutrophils, has been observed (22). These findings support our hypothesis of a predominantly sterile inflammatory response in the dental pulp due to hypomineralized enamel (57). Nevertheless, we cannot exclude more subtle immunological changes, including shifts within the subpopulations of the resident macrophages (58), alterations in other myeloid cells, or shifts in adaptive immunity, especially considering the low-grade chronic stimuli that the dental pulp receives (59, 60).

Soft and porous enamel not only elevates the transient activation of the sensory organ, but these alterations appear to place the dentin-pulp complex in a *primed* state of heightened sensory and odontoblastic responsiveness. This is consistent with clinical observations of persistent hypersensitivity and, reduced anesthetic efficacy during dental treatments, and reduced dental pulp oxygen saturation (61–63), in patients with molar hypomineralization, which are likely to be consequences of inflammatory changes in the dentin-pulp complex and an altered sensitized state of the sensory organ of the tooth due to chronic stimuli (16, 64, 65). Surprisingly, we found that this *primed* state in teeth with hypomineralized enamel, protected the dental pulp against greater insults; with reduced sensory afferent loss after pulp injury, compared to teeth with intact enamel. Whether this reflects increased resistance (enhanced inflammatory response) or adaptation (tolerance to sustained stress) remains unclear (34, 66). Furthermore, the relative contribution of sensory neurons (67), odontoblasts, and the immune cells (68) to this protective response requires further mechanistic studies.

The observed protective effects and tissue responses due to priming of the dentin-pulp complex should be interpreted with caution from a clinical perspective. While the pulp beneath hypomineralized enamel may be more resilient to acute injury, it does not mitigate the well-established increased risk of caries associated with hypomineralized enamel (69–71). Nevertheless, emerging clinical evidence suggests that pulpal healing capacity in hypomineralized teeth may be comparable to that of unaffected teeth (72).

Several limitations should be acknowledged. First, our use of a global KLK4 KO model may cause off-target effects; however, KLK4 expression in other tissues is very limited and should not interfere with our findings (73, 74). However, validation in conditional knock-out models will be important in future studies. Second, the current study does not address the temporal dynamics of these responses following tooth eruption. Enamel hypomineralization is a developmental defect, and exposure to increased stimuli starts as early as tooth eruption. It is important to understand how the sensory nervous system, at the peripheral and central level, adapts to chronic stimulation throughout the lifespan of the teeth with enamel defects, such as molar hypomineralization (75). Therefore, longitudinal studies will be essential to understand how sensory, cellular, and immune responses in the dentin-pulp complex evolve over time, including any post-eruptive changes in enamel (76).

## MATERIALS & METHODS

### Animals

All animal experiments are approved by the Institutional Animal Care and Use Committees (IACUC) (HSDM IS00002576-3, HMS IS00000054-9, AFI 25-004). The mice used were housed and cared for by qualified veterinarians and veterinary technicians and monitored for pain and postoperative distress in accordance with IACUC standards. Experiments were performed using age-matched male and female mice. The animals used were 6-8 weeks old at the time of sacrifice. The mouse lines used in the present study included C57BL/6J and KLK4 KO (generously provided by Dr. Hu and Dr. Simmer to Dr. Bidlack).

### Tissue Collection

#### Oral swabs

For oral microbiome analysis, five mice per group were used. The previously published protocol was followed for the collection (21). Briefly, following isoflurane anesthesia, mice were scruffed and held in a position that allowed the placement of the ultra-fine polyester-tipped swab in the oral cavity. The oral mucosa facing the teeth and the tooth surfaces were swabbed for one minute. Each swab was individually placed in a microcentrifuge tube containing an RNA stabilization solution (RNA-DNA Shield, Zymo Research Corporation, CA).

#### Dental pulp cells

Dental pulp cell suspension was prepared as previously described(22). Briefly, dissected maxillae and mandibles were placed in minimum essential medium (MEMα, Gibco, Thermo Fisher Scientific, MA). Only the crowns of molars were collected by cutting them at the crown-root interface. Enzymatic digestion (4 mg/ml collagenase and 4 mg/ml dispase II) of the pulp tissue in molar crowns was performed at 37 °C for 1 hour. The pulp tissue was detached from the dentin by vigorous pipetting and then filtered through a 70 μm mesh. The samples were spun down at 300g for 5 minutes and washed with flow buffer (2% FBS, 1 mM EDTA, 0.1% sodium azide).

#### Maxillae, incisor teeth, trigeminal ganglia and brainstem

Animals were perfused with PBS containing heparin, followed by 4% paraformaldehyde (PFA). All tissues were post-fixed with 4% PFA overnight. Maxillae were first decalcified using 0.5 M EDTA at 4°C for 4 weeks. For frozen samples, tissues were processed with 30% sucrose for 2 days, then with 50% sucrose for 2 days at 4°C and finally cryopreserved using an optimal cutting temperature compound (OCT). Maxillae and brainstem were sectioned at 30 μm, and trigeminal ganglia were sectioned at 14 μm. For paraffin embedding, tissue processing was performed using a tissue processor (TP1020, Leica). Maxillae were embedded with paraffin and sectioned (7 μm). Mandibular incisors sectioned at the bone level were also collected from the same animals and were used to grow biofilm on the buccal surface.

### Ex vivo Biofilm Growth

Dissected mandibular incisors were disinfected in a 2.5% NaOCl solution for 5 minutes, then were ultrasonicated in sterile water twice for 5 minutes. The incisors were then placed in individual wells in a 48-well plate to grow a single species biofilm on the buccal surface. Pellicle formation was simulated by incubating the enamel surface with 500 μl of filtered (0.22 μm) human saliva for one hour. Then the saliva was removed and *Streptococcus mutans (*UA140) was grown at 5% CO_2_ and 37°C using 500μl of BHI (Brain Heart Infusion) medium. After 48 hours, the incisors were rinsed with sterile PBS to remove non-adherent cells, and the biofilm was fixed using 50% ethanol. Finally, incisors with biofilm on their surfaces were transferred to custom-made sample holders and processed for fluorescent in situ hybridization (FISH) and imaged with a confocal microscope as described under Microscopy Imaging.

### Flow Cytometry

Single pulp cells were resuspended in flow buffer (PBS with 2% FBS and 1 mM EDTA). After Fc receptor blockade for 10 minutes on ice, cells were washed and incubated for 30 minutes at 4°C with the following surface antibodies: Zombie Aqua Fixable Viability Dye (1:1000, BioLegend), CD45-APC/Cy7 (1:100, BioLegend), CD11b-PE (1:100, BioLegend), F4/80-PB (1:100, BioLegend), CX3CR1-FITC (1:100, BioLegend), and Ly6G-APC (1:100, BioLegend). Following staining, cells were washed, fixed in 1% paraformaldehyde, and filtered through a 40 μm mesh. Flow cytometry was performed using a CytoFLEX flow cytometer (Beckman Coulter Life Sciences), and data were analyzed with FlowJo software (FlowJo LLC). Gating strategy excluded doublets and dead cells, and immune subsets were identified as follows: CD45+ immune cells, CD11b⁺ myeloid cells, F4/80⁺ macrophages, CX3CR1⁺/⁻ macrophage subsets, and CD11b⁺F4/80⁻Ly6G⁺ neutrophils.

### 16S RNA DNA Extraction, Sequencing and Bioinformatics

Oral swabs were collected and immersed in RNA-DNA Shield solution. Samples were then submitted to Forsyth Oral Microbiome Core (FOMC) for DNA extraction and 16S rRNA sequencing. DNA extraction was performed using ZymoBIOMICS 96 Magbead DNA Kit, and library preparation and next-generation sequencing were performed at Zymo Research (Irvine, CA). Quick-16S™ NGS Library Prep Kit (Zymo Research, Irvine, CA) was used for targeted sequencing of the DNA samples. Quick-16S™ Primer Set V1-V3 (Zymo Research, Irvine, CA) was used to prepare custom-designed primers that provided the best 16S gene coverage with high sensitivity. The final library was sequenced on Illumina® MiSeq™ with a V3 reagent kit (600 cycles). The sequencing was performed with 10% PhiX spike-in.

Raw sequence quality control, noise filtering, pair reads merging, and chimera filtering were performed at FOMC using the DADA2 pipeline, as well as taxonomy assignment, alpha and beta diversity analyses, and differential abundance analysis. Shannon alpha diversity differences were calculated with the Kruskal-Wallis H test. For analysis of beta diversity, a Bray-Curtis dissimilarity matrix was calculated and plotted by the non-metric multi-dimensional scaling (NMDS).

### Trigeminal Ganglion RNA Isolation, Library Construction and Sequencing

Dissected trigeminal ganglia were snap-frozen in liquid nitrogen and shipped on dry ice to Zymo Research (Irvine, CA). RNA extraction was performed using the Quick-RNA Microprep Kit, and 3’ mRNA-Seq libraries were constructed from total RNA. To enrich for poly(A) RNA molecules, an oligo(dT) primer with a partial P5 adapter was used. After that, reverse transcription was performed, followed by partial P7 adapter ligation and second-strand cDNA synthesis. Lastly, libraries were amplified to incorporate full-length adapters. Successful library construction was confirmed with Agilent’s D1000 ScreenTape Assay on TapeStation. mRNA-Seq libraries were sequenced on an Illumina NovaSeq to a sequencing depth of at least 30 million read pairs (150 bp paired-end sequencing) per sample.

The bioinformatics pipeline is based on the nf-core/rnaseq pipeline (23). The Deseq2 package on R was used for the differential expression analysis. Further statistical analysis of the data is described under the Statistical Analysis section.

### MicroCT Analysis

Dissected maxillae of WT and KLK4 KO were analyzed by microcomputed tomography(microCT) on a Scanco µCT40 scanner (Scanco Medical, Switzerland) with a 6 μm voxel resolution. Grayscale images were obtained from a virtual slice that goes through all three molar teeth to demonstrate the mineral density difference in WT and KLK4 KOs.

### Pulp Injury Procedure

Mice were anesthetized via intraperitoneal injection of 100 mg/kg ketamine + 10 mg/kg xylazine in sterile phosphate-buffered saline (PBS). The exposures were generated according to published protocols (22) . Briefly, the maxillary first molar tooth was drilled with a ¼ round bur until the middle pulp horn was mechanically exposed and left open. The contralateral molar served as the uninjured control.

### Tissue Processing for Histology

#### Immunofluorescence Staining

Cryosections were first dried for 1 hour and washed with PBS, and then blocked with 5% goat serum (Sigma-Aldrich) and 1% Triton X-100 in PBS (0.25% Triton X-100 for the trigeminal ganglion immunostaining for ATF3) for 1 hour at room temperature. Overnight incubation at 4 °C with the primary antibodies was performed. The antibodies used were rabbit anti-βIII tubulin (Tuj1, 1:500, Abcam), rat anti-Ly6G/C (1:200, Abcam), rabbit anti-CD31 (1:150, Novus Biologicals), rabbit anti-cFos (1:1000, Cell Signaling Technology), rabbit anti-ATF3 (1:200, Sigma-Aldrich). Sections were washed, followed by incubation for 2 hours at room temperature with secondary antibodies. Secondary antibodies were: goat anti-rabbit Alexa fluor 546 (1:500, ThermoFisher Scientific), goat anti-rat Alexa fluor 647 (1:500, ThermoFisher Scientific). Sections were mounted using an antifade mounting medium with DAPI (Vector Laboratories). Image acquisition was performed using a Leica Stellaris confocal microscope or a Leica THUNDER DMi8 Inverted Autofocus fluorescence microscope. Image acquisition in experiments was conducted under identical exposure and acquisition settings across samples. Image processing and quantification were performed in Fiji (ImageJ) using standardized parameters.

#### Hematoxylin and eosin (H&E) Staining

After embedding in paraffin and sectioning, samples were rehydrated, stained with H&E, dehydrated, and mounted (24). Coronal pulp in the sections was examined by a board-certified oral and maxillofacial pathologist, blinded to experimental groups (L.A.), as previously performed (8). Briefly, coronal pulp was scored for the extent of necrosis (0-3; 0: no necrosis, 1: 1/3 of the area of interest is necrotic, 2: 2/3 of the area of interest is necrotic, 3:3/3 of the area is necrotic) and the extent of odontoblast loss (0-3; 0: no changes, 1: 1/3 of the area of interest is affected, 2: 2/3 of the area of interest is affected, 3:3/3 of the area is affected). The presence of edema is scored as binary (0: absent; 1: present). Three sections per animal were scored and averaged to obtain the individual animal score.

#### Fluorescence In Situ Hybridization (FISH)

The biofilm grown on incisor surfaces was labeled **with** the oligonucleotide probe Str405-AT488 (biomers.net, Germany) targeting 16S rRNA. FISH was performed at 46°C for 4 hours using a hybridization buffer (2pmol/ul probe, 900mM NaCl, 20mM Tris HCl, 0.01%SDS, 20% HiDi) (25). Following hybridization, the incisors were rinsed with a wash buffer (215 mM NaCl, 20 mM Tris-HCl, 5 mM EDTA). Finally, incisors were mounted for imaging (Prolong Antifade Medium, Thermofisher Scientific, MA).

### Microscopy Imaging & Quantifications

#### Reactionary Dentin Area

Using the phase contrast images from the maxillary molars, reactionary dentin was manually selected based on its irregular tubular structure, which is distinct from the primary and secondary dentin (Figure 1A, yellow dashed line). Reactionary dentin area was quantified separately for each cusp (mesial, mid, distal), and was normalized to the total pulp chamber area of each tooth.

#### Sensory afferent withdrawal distance in the dental pulp

Molar teeth sections stained for anti-βIII tubulin, were processed in Fiji to mark the outermost layer of Raschkow Plexus and the innermost layer of the dentin (Figure 2C, yellow arrowheads). A Fiji plugin was then used to measure the distance between these lines at 20-pixel intervals. Mean values of this distance were then calculated for each cusp.

#### ATF3 in trigeminal ganglia

Four animals (three sections per animal) per group were included to quantify ATF3 at the intersection area between V3 and V2 divisions of the trigeminal ganglion. Images were quantified in Fiji using the “Analyze Particles” function. The number of ATF3-positive cells was quantified and normalized to the area of the trigeminal ganglion section.

#### cFos in trigeminal nuclei

Four animals per group were used to image three sections of the caudalis and interpolaris regions of the trigeminal nucleus per animal. Two independent experiments were performed. Images were processed in Fiji to distinguish c Fos-positive nuclei within the location of the trigeminal nucleus. Regions were identified based on the Allen Mouse Brain Atlas(26). The number of positive nuclei was quantified using the “Analyze Particles” function. The co-localization of c-FOS signals with DAPI was reviewed, and the % area with positive cells was calculated.

#### Afferent Loss

Quantification of afferent loss was performed in Fiji by measuring the Tuj1-negative area within the pulp and expressing it as a percentage of the total pulp area.

#### Vascularization

Vascularization was quantified using a custom Fiji macro adapted from Pinho-Ribeiro(27) to calculate the total length of CD31⁺ vascular segments, normalized to the total selected pulp area.

#### Biofilm Coverage

Biofilm on the incisor surface was imaged using an LSM 980 confocal microscope (Zeiss, US). 5 pairs of mandibular incisors were imaged for each group and the Str405-AT488 oligonucleotide was excited with a 488nm laser. 15 virtual slices were obtained from each incisor using 1AU settings. Biofilm coverage area was quantified on maximum-intensity projections of a z-stack consisting of 15 sections at 1AU using FIJI. Quantifications were normalized to the total surface area of each incisor.

### Statistical Analysis

An unpaired Student’s t-test was used to compare two groups for a single variable with normal distributions. Unpaired t-test or two-way ANOVA with Sidak’s or Tukey’s multiple comparisons tests were performed when appropriate. A p-value less than 0.05 was considered statistically significant. Statistical analyses and data visualization were performed in Prism 9 (GraphPad Software, US).

Along with GeneIDs, adjusted p-values, and log2 (fold change) from the trigeminal ganglion RNA sequencing analysis, the data were processed in R using the readxl and dplyr packages to remove duplicates and further format the data. The volcano plot was prepared with the enhancedvolcano package, using 0.05 as the adjusted p-value cutoff, and 1 as the fold change cutoff (28). Finally, the same dataset was used for the GO Enrichment analysis, which was performed using the topGO package on R (29).

## Data Availability

Source data presented in the figures are provided in Supplementary Data 1. Raw sequencing data used in this study have been deposited in the NIH NCBI Sequence Read Archive (SRA) with the BioProject code PRJNA1513133 and is available at https://dataview.ncbi.nlm.nih.gov/object/PRJNA1513133?reviewer=i9v1k286gtuppgne75jb73uh7p. Any other data/information is available upon request.

## Acknowledgements

H.K. discloses support for this work from NIH/NIDCR [K08DE033793], O.E. discloses support for this work from NIH/NIDCR [K99DE033421, R00DE033421]. JLG and FBB acknowledge support for this work from the Forsyth-HSDM Pilot Grant. Schematics in the figures were prepared using biorender.com. We would like to thank all members of the Chiu Lab at Harvard Medical School for their generous help and feedback. Confocal microscopy was performed at the ADA Forsyth Institute Advanced Microscopy Core Facility (RRID: SCR_021121).

## Author Contributions

H.K. contributed to conception, data acquisition, and analysis and interpretation, and drafted the manuscript; R.B. and M.S. contributed to data acquisition and analysis; L.A. contributed to data analysis and interpretation of the data, E.W. contributed to drafting the manuscript review. X.H. contributed to the interpretation of the data and manuscript. I.M.C., F.B.B., and J.L.G. contributed to the conception, interpretation of the data, and manuscript review and revision O.E. contributed to conception, design, data acquisition, and analysis, writing and revision of manuscript.

## Competing Interests

The authors declare no competing financial or non-financial interests.

## Notes

https://dataview.ncbi.nlm.nih.gov/object/PRJNA1513133?reviewer=i9v1k286gtuppgne75jb73uh7p

### Competing Interest Statement

The authors have declared no competing interest.

